# Thor-Ribo-Seq: ribosome profiling tailored for low input with RNA-dependent RNA amplification

**DOI:** 10.1101/2023.01.15.524129

**Authors:** Mari Mito, Yuichi Shichino, Shintaro Iwasaki

## Abstract

Translation regulation plays a pivotal role in the diversification of gene expression and the response to intra- and extracellular environmental cues. Ribosome profiling (or Ribo-Seq) serves as a sensitive, quantitative, comprehensive, and data-rich technique to survey ribosome traversal across the cellular transcriptome. However, due to the intricacy of library preparation, applications to low input have presented analytic challenges. To overcome this issue, here we developed Thor-Ribo-Seq, a ribosome profiling method tailored for low input. Thor-Ribo-Seq harnesses RNA-templated RNA transcription to linearly amplify ribosome footprints, assessing ribosome traversal at codon resolution with limited artifacts. This highly sensitized ribosome profiling approach provides a versatile option to investigate the translatome in precious samples.

## Introduction

Although the transcriptomic profile has been recognized as a proxy for gene expression, recent studies have shown that regulation at the level of protein synthesis has a profound effect on the final output of genes. Ribosome profiling (or Ribo-Seq), a technique based on RNase footprinting of mRNAs by ribosomes and subsequent deep sequencing, has emerged as a game-changing approach to study protein synthesis in cells^1^. The applications of this technique are highly manifold; it can measure translation efficiencies across the transcriptome^2,3^, assign open reading frames (ORFs) *de novo*^4^, explore ribosome movement speed at codon resolution^2,3^, and even estimate the structural status of ribosomes^5,6^. The widespread applications of this technique in diverse materials and biological conditions have unveiled previously uncharacterized layers of regulation of protein synthesis^2–4^. The road ahead of ribosome profiling/Ribo-Seq should involve application to rare, precious, and difficult-to-prepare samples. However, standard protocols require a significant amount of cells for high-quality library preparation^7,8^ and thus present technical hurdles for low inputs.

Therefore, efforts to optimize ribosome profiling/Ribo-Seq with small amounts of materials are underway. A one-pot reaction for library preparation is an advantageous approach to avoid sample loss. For this purpose, a ligation-free method could be applied to isolated ribosome footprints; this system allows the sequential reactions of 3’ end poly(A) tailing, reverse transcription, and linker addition by template switching in the same tube^9–12^. The recently reported technique of single-cell ribosome profiling (scRibo-Seq) started the one-pot reaction with the cell lysate, conducting RNase digestion, first linker ligation, reverse transcription, second linker ligation, and PCR^13^. However, these approaches do not have any means to amplify the DNA or RNA before PCR and thus involve the risk of material loss during the procedures.

RNA amplification with T7 RNA polymerase-mediated *in vitro* transcription has been a useful option to handle low amounts of nucleic acids^14,15^. Because of the linear amplification, this strategy has been reported to be a less biased approach^16^ than exponential expansion by PCR. In practice, linear amplification has been implemented in diverse single-cell sequencing techniques, such as whole-genome analysis^17^, RNA-Seq^18,19^, and epigenomic profiling^20,21^. Similarly, the investigation of limited transcriptomes on RNA binding proteins was greatly improved by this approach^22,23^. However, all those methods required the formation of double-stranded DNA (dsDNA) for *in vitro* transcription; thus, the execution of this technique was possible only at later steps of library preparation. An amplification strategy before dsDNA synthesis should be beneficial for sequencing techniques targeting RNA because its execution at an early step of the procedures effectively counteracts material loss.

Here, we developed Thor-Ribo-Seq tailored for low material input. This technique harnesses RNA-dependent RNA amplification by T7 RNA polymerase, which can use an RNA-DNA chimera as an *in vitro* transcription template. Rationally designed linker oligonucleotides maximize the linear expansion of material and allow the computational suppression of bias. Ultimately, Thor-Ribo-Seq provides a widely applicable platform for translation study with ultralow and even standard inputs.

## Results

### Implementation of RNA-dependent RNA amplification into Ribo-Seq

An apparent pitfall of Ribo-Seq is the necessary material amount or cell number for library preparation. Indeed, standard ribosome profiling experiments (Figure S1A) typically required 1-10 μg of total RNA, which corresponds to approximately 10^5^-10^6^ cells of human embryonic kidney (HEK) 293. Lower quantities of cell lysate, such as extract corresponding to 0.1 μg of total RNA (or ~10^4^ cells), could not enable DNA library amplification by PCR (Figure S1C) due to sample loss during the multiple complicated steps in library generation. This drawback ultimately restricted ribosome profiling experiments for limited samples.

To overcome this issue, we applied the Thor (T7 High-resolution Original RNA) strategy (implemented in the LUTHOR 3’ mRNA-Seq Library Prep Kit provided by Lexogen) to amplify the linker-ligated ribosome footprints at an early stage of library preparation (Figures 1A and S1B) before loss of material. We designed the linker DNA oligonucleotide with the antisense sequence of the T7 promoter. Hybridization of short DNA oligonucleotides covering the antisense sequence created a partial dsDNA region competent for RNA transcription^24^. Since T7 polymerase can synthesize complementary RNA from the RNA template conjugated with the dsDNA T7 promoter region^25^, this configuration allows amplification of the complementary RNA of ribosome footprints. We also employed the optimized DNA sequence upstream and downstream of the T7 promoter to maximize transcription^26^. For multiplexing, the linker also has a sample barcode sequence (Figure S1B). To suppress the amplification bias, we added a unique molecular index (UMI) in the linkers (Figure S1B) (see below for details).

**Figure 1.**
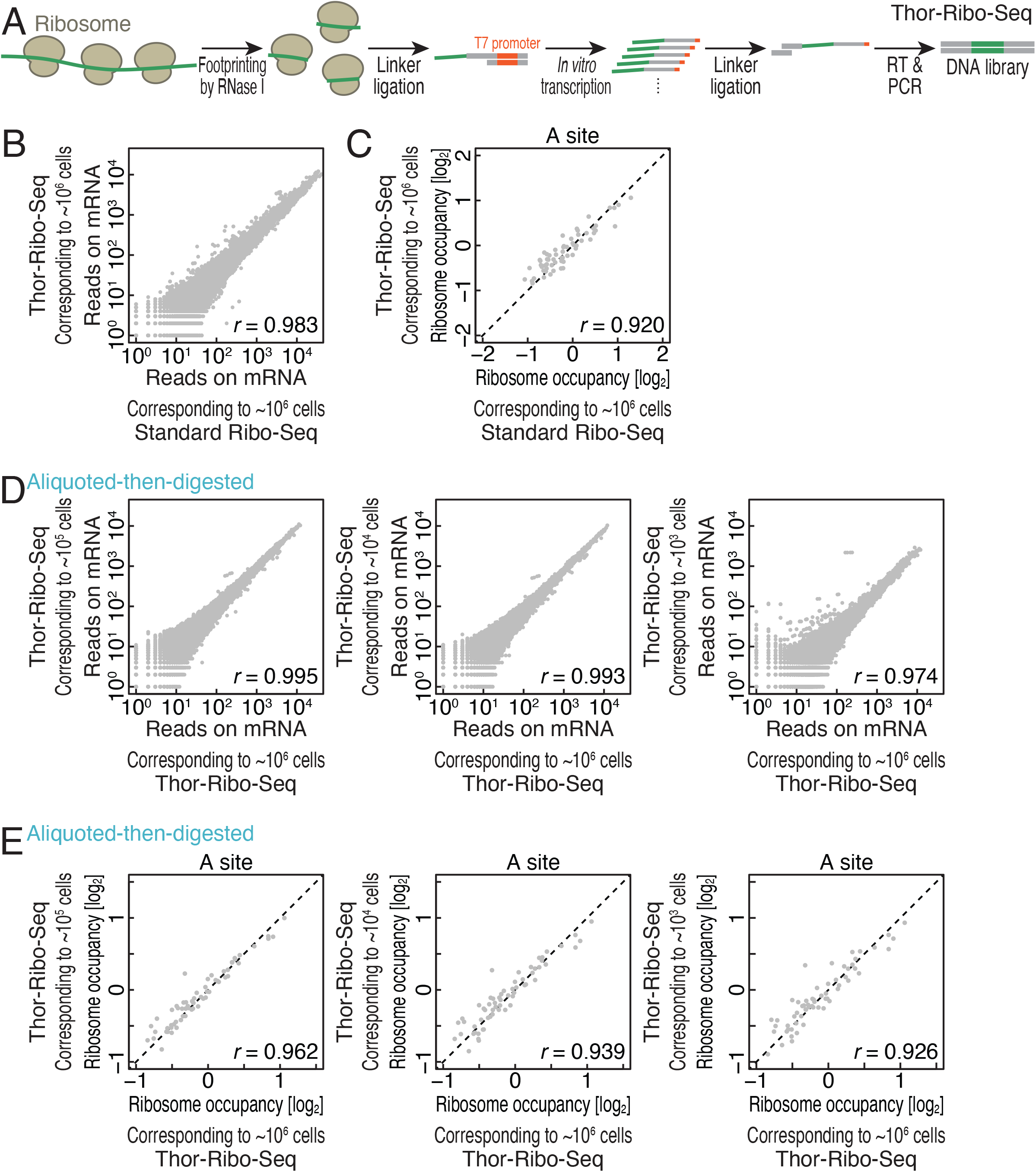
Thor-Ribo-Seq assesses translation status from low inputs. (A) Schematic representation of the library preparation strategy of Thor-Ribo-Seq. RT, reverse transcription. (B) Correspondence of reads on transcripts between standard Ribo-Seq and Thor-Ribo-Seq with normal inputs. (C) Correspondence of averaged ribosome occupancy on A-site codon sequences between standard Ribo-Seq data and Thor-Ribo-Seq with normal inputs. (D) Correspondence of reads on transcripts across Thor-Ribo-Seq with a normal input and that of “aliquoted-then-digested” experiments with low material inputs. (E) Correspondence of averaged ribosome occupancy on A-site codon sequences across Thor-Ribo-Seq with a normal input and that of “aliquoted-then-digested” experiments with low material inputs. *r*, Pearson’s correlation coefficient. See also Figures S1, S2, S3, and S4.

We set out to compare the performance of the “Thor”-Ribo-Seq with standard Ribo-Seq with the normal amount of material (10 μg of total RNA or ~10^6^ cells). We observed hallmarks of ribosome footprints in Thor-Ribo-Seq: read peaks at 22 and 28 nucleotides (nt) (Figure S2A) and strong triplet periodicity along the CDS (Figure S2B-C) as in standard Ribo-Seq. Also, both Ribo-Seq methods detected comparable numbers of genes (Figure S2D). Thus, high correspondence of the reads on ORFs was found between standard Ribo-Seq and Thor-Ribo-Seq (Figure 1B).

Moreover, Thor-Ribo-Seq exhibited limited artifacts in codon-wise examinations of ribosome occupancy. A-site codon sequences are the most prominent determinants of ribosome duration time at each codon^27–36^ and thus indicated differential ribosome occupancies (Figure 1C). The ribosome occupancy divergence was consistently maintained in Thor-Ribo-Seq (Figure 1C). These results indicate that Thor-Ribo-Seq strategy provides a bias-less option at least for a conventional amount of samples.

### Thor-Ribo-Seq for translational profiling from low material input

Strikingly, Thor-Ribo-Seq strategy allowed us to perform ribosome profiling with inputs too low to be processed by standard Ribo-Seq. Given that RNase treatment^10,37–39^ and the downstream processes^29,33^ introduce different biases, we designed experiments to control the two impactful sources individually when Thor-Ribo-Seq is applied to small inputs. To maintain the same RNase treatment conditions, we prepared a library with aliquots of RNase-treated lysate (denoted as “digested-then-aliquoted”) (Figure S3A); we treated cell lysate containing 10 μg of total RNA (or ~10^6^ cells) with RNase I, as in standard ribosome profiling, and then divided the reaction into aliquots corresponding to 0.01, 0.1, or 1 μg of total RNA for downstream Thor-Ribo-Seq library preparation. Transcription by T7 polymerase enabled the amplification of the complementary RNA (Figure S3B) and thus the sequencing library (Figure S3C), for inputs at least as low as 0.01 μg (equal to ~10^3^ cells). In the sequencing data, the benchmarks of footprints, including read length (Figure S4A) and 3-nt periodicity (Figure S4B-D), were maintained.

We assessed translation status across the transcriptome in the “digested-then-aliquoted” Thor-Ribo-Seq experiments. The reads on ORFs exhibited high correlations to the Thor-Ribo-Seq data prepared with ~10^6^ cells (Figure S4I). Moreover, our improved method yielded ribosome footprints from a large number of genes (Figure S4J), suggesting only a marginal limitation of the detection sensitivity. The slight reduction at the lowest input (~10^4^ cells) may originate from the shallow complexity of the transcripts in the lysate. In addition to the mRNA-wise analysis, our approach allowed codon-wise examinations of ribosome occupancy from low inputs (Figure S4K). These data indicated that if any biases stem from the steps after RNase digestion, they do not hamper Thor-Ribo-Seq translatome analysis for small inputs.

Then, we conducted the whole Thor-Ribo-Seq procedure, including RNase digestion and downstream library preparation from the low material inputs (denoted as “aliquoted-then-digested”) and assessed the bias derived from all steps (Figure S1A). As observed in the “digested-then-aliquoted” condition, the libraries were successfully prepared, irrespective of the material amounts (Figure S3B and S3C). The entire properties of the Ribo-Seq—footprint size, 3-nt periodicity, and gene detection—were remarkably consistent in these experiments, even with the lowest inputs (equal to ~10^3^ cells) (Figure S4E-J). Ultimately, “aliquoted-then-digested” Thor-Ribo-Seq also showed high correspondence to the Thor-Ribo-Seq data from ~10^6^ cells in the mRNA-wise analysis (reads on transcripts) (Figure 1D and S4I) and in codon-wise ribosome occupancy (Figure 1E).

We noted that the suppression of read duplications generated by T7 polymerase-mediated RNA amplification benefitted the adequate evaluation of data. Our Thor-Ribo-Seq library contained unique molecular indexes (UMIs) at two distinct positions: at the 3’ end of footprints introduced by the first linker and at the 5’ end of footprints originating from the second linker (Figures S1B and S4L). Given the timing of the UMI addition, 3’ end UMI suppressed the read duplications that occurred in the *in vitro* transcription, and the combination of 3’ and 5’ UMIs limited those duplications in the PCR (Figure S4L). We tested the potency of the UMIs to restrain the biases and found that the minimization of overamplified reads by *in vitro* transcription led to a high correlation among Thor-Ribo-Seq data compared to the same correction of biases in PCR (Figure S4I and S4M). Our analyses mentioned above used 3’ UMI suppression unless noted.

## Discussion

This study developed Thor-Ribo-Seq as a sensitive methodology to perform translatome analysis with limited material, maintaining the characteristic features of ribosome profiling experiments. Our strategy outperforms in A-site offset assignments compared to preexisting methods for small inputs. Micrococcal nuclease (MNase), which has been used for scRibo-Seq^13^ due to the stringent activity control by Ca^2+^ ions, has an A/U preference. The poly(A) tailing and template switching used in the one-pot reaction^9–12^ add homopolymeric nucleotides to the footprints and thus generate ill-defined read boundaries at both the 5’ and 3’ ends. These drawbacks hamper the A-site offset estimation in ribosome footprints. In contrast, our strategy uses less biased RNase I and double-linker ligation to explicitly determine the ends of reads, preventing ambiguity regarding the A-site position reference.

Although our method requires custom-made linkers, all the other reagents are commercially available. Thus, this technique should be easy to implement in library construction not only for regular ribosome profiling but also for selective ribosome profiling^40,41^, translation complex profile sequencing (TCP-Seq or 40S footprinting)^42,43^, selective TCP-Seq^44–47^, and cross-linking and immunoprecipitation (CLIP)-Seq^48^, all of which handle limited amounts of RNA fragments for deep sequencing.

## Acknowledgments

We are grateful to all the members of the Iwasaki laboratory for constructive discussions, technical help, and critical reading of the manuscript. We also thank the HOKUSAI SailingShip supercomputer facility at RIKEN for computation support. S.I. was supported by the Ministry of Education, Culture, Sports, Science and Technology (MEXT) (a Grantin-Aid for Transformative Research Areas [B] “Parametric Translation”, JP20H05784), AMED (AMED-CREST, JP22gm1410001), and RIKEN (“Biology of Intracellular Environments” and “Integrated life science research to challenge super aging society”). Y.S. was supported by JSPS (a Grant-in-Aid for Early-Career Scientists, JP21K15023), MEXT (a Grant-in-Aid for Transformative Research Areas [A] “Multifaceted Proteins”, JP21H05734), and RIKEN (Special Postdoctoral Researchers and Incentive Research Projects).

## Author contributions

Conceptualization, M.M., Y.S., and S.I.

Methodology, M.M., Y.S., and S.I.

Formal analysis, M.M. and Y.S.

Investigation, M.M. and Y.S.

Writing – Original Draft, Y.S. and S.I.

Writing – Review & Editing, M.M., Y.S., and S.I.

Visualization, Y.S.

Supervision, Y.S. and S.I.

Project Administration, S.I.

Funding Acquisition, Y.S. and S.I.

## Experimental procedures

### Cell lines

Flp-In T-REx 293 cells (Thermo Fisher Scientific, R78007) were cultured in DMEM, high glucose, GlutaMAX Supplement (Thermo Fisher Scientific) with 10% FBS at 37°C with 5% CO2, according to the manufacturer’s instructions. The cell culture was routinely tested for *Mycoplasma* contamination with the e-Myco VALiD Mycoplasma PCR Detection Kit (iNtRON Biotechnology) and confirmed to be negative.

### Library construction for Ribo-Seq

#### Cell lysate preparation

The cell lysate was prepared as described previously^8^. Briefly, Flp-In T-REx 293 cells cultured in a 10-cm dish were washed with ice-cold PBS and then lysed with lysis buffer [20 mM Tris-HCl pH 7.5, 150 mM NaCl, 5 mM MgCl_2_, 1 mM dithiothreitol (DTT), 1% Triton X-100, 100 μg/ml cycloheximide, and 100 μg/ml chloramphenicol]. The extract was treated with 0.025 U/μl Turbo DNase (Thermo Fisher Scientific), clarified by centrifugation at 20,000 × g at 4°C for 10 min, flash-frozen in liquid nitrogen, and stored at −80°C. The RNA concentration in the lysate was measured by a Qubit RNA BR Assay Kit (Thermo Fisher Scientific). Of note, cells were treated with DMSO for 15 min in the standard Ribo-Seq.

#### Standard Ribo-Seq

Standard Ribo-Seq was conducted as described previously^8^. Cell lysate containing 10 μg of RNA was brought to a volume of 300 μl by lysis buffer, treated with 20 U of RNase I (LGC Biosearch Technologies) (*i.e*., 0.067 U/μl) at 25°C for 45 min, placed on ice, supplemented with 200 U of SUPERase•In RNase Inhibitor (Thermo Fisher Scientific), and ultracentrifuged in a sucrose cushion at 100,000 rpm at 4°C for 1 h with a TLA110 rotor (Beckman Coulter). The ribosome pellet was dissolved in TRIzol reagent (Thermo Fisher Scientific), and RNA was isolated with Direct-zol RNA MicroPrep (Zymo Research). The RNA fragments corresponding to 17-34 nt in length were excised from the gel, dephosphorylated by T4 polynucleotide kinase (New England Biolabs), ligated to linkers by T4 RNA Ligase 2, truncated KQ (New England Biolabs), rRNA-depleted by a Ribo-Zero Gold rRNA Removal Kit (Human/Mouse/Rat) (Illumina), and reverse transcribed by ProtoScript II Reverse Transcriptase (New England Biolabs). Then, the cDNA was circularized by CircLigaseII ssDNA ligase (LGC Biosearch Technologies) and amplified by PCR with Phusion High-Fidelity DNA Polymerase (New England Biolabs). DNA libraries were sequenced on HiSeq 4000 (Illumina) with a single-end 50 bp option.

#### Thor-Ribo-Seq

For the “aliquoted-then-digested” experiments, cell lysates corresponding to 1, 0.1, and 0.01 μg of RNA were scaled up to 300 μl by lysis buffer, digested with 0.067 U/μl RNase I at 25°C for 45 min, placed on ice, supplemented with 200 U of SUPERase•In RNase Inhibitor, and subjected to a sucrose cushion.

For the “digested-then-aliquoted” experiments, cell lysates were treated with RNase I and SUPERase•In RNase Inhibitor as described in “Standard Ribo-Seq”. Then, aliquots corresponding to 1, 0.1, and 0.01 μg of RNA were brought up to a volume of 300 μl by lysis buffer and subjected to a sucrose cushion.

RNA isolation and gel excision were performed as described previously^8^. The first linker (SI181) 5’-/Phos/NNNNN**ATCGT**AGATCGGAAGAGCACACGTCTGAACTCCAGTCA*CCCT ATAGTGAGTCGTATTAGTCA/ddC/-3’*, where /Phos/ represents a 5’ phosphate, /ddC/ represents a terminal 2’, 3’-dideoxycytidine, N represents a random nucleotide for UMI, bold characters represent a sample barcode sequence, and italicized characters correspond to the T7 promoter, and the second linker SI183 5’-/Phos/NNAGATCGGAAGAGCGTCGTGTAGGGAAAGAG/ddC/-3’ was preadenylated by Mth RNA Ligase (New England Biolabs) as described previously^8^. After dephosphorylation by T4 polynucleotide kinase, RNAs were ligated with the preadenylated first linker SI181 by T4 RNA Ligase 2, truncated KQ. After hybridization of the T7 promoter oligonucleotide SI182 5’-TGAC*TAATACGACTCACTATAGGG*TGACTGGAGTTCAG-3’, complementary RNA was transcribed by a T7-Scribe Standard RNA IVT Kit (CELLSCRIPT) at 37°C for 2.5 h and gel-purified. After dephosphorylation by T4 polynucleotide kinase, RNAs were ligated to the preadenylated second linker SI183 by T4 RNA Ligase 2, truncated KQ, hybridized to SI186 5’-/Phos/AATGATACGGCGACCACCGAGATCTACACTCTTTCCCTACACGACGCTC-3’, and then reverse-transcribed by ProtoScript II Reverse Transcriptase. cDNA was PCR-amplified with NI798 and NI799 5’-CAAGCAGAAGACGGCATACGAGATCGTGATGTGACTGGAGTTCAGACGTGTG-3’ by Phusion High-Fidelity DNA Polymerase. rRNA depletion by the Ribo-Zero unit from TruSeq Stranded Total RNA Library Prep Gold was applied to cDNAs but can be used for the first linker-ligated RNAs.

As an alternative to the first linker and the hybridizing oligonucleotide, SI96_001 5’-/Phos/NNNNN**CATATTCCTGGTGG**AGATCGGAAGAGCACACGTCTGAACTC CAGTCACATTAT*CCCT4T4GTGHGTCGT4TZ4*AATTC/SpC3/, where /SpC3/indicates the 3’ C3 spacer, and SI194 5’-GAA*TTTAATACGACTCACTATAGGG*ATAATGTGACTGGAGTTCAG-3’ could be used.

DNA libraries were sequenced on HiSeq X (Illumina) with a paired-end 150-bp option.

### Deep sequencing data analysis

Data processing was performed as previously described^7^ with modifications. Using Fastp v0.21.0^49^, bases of pair-end reads were corrected by overlap analysis, and then quality filtering and removal of the adapter sequence (AGATCGGAAGAGCACACGTCTGAA) were performed on read 1. For single-end reads, base correction was skipped. After the extraction of UMI and barcode sequences on the linkers by a custom script (fastx-split in https://github.com/ingolia-lab/RiboSeq), reads mapped to noncoding RNAs using STAR v2.7.0a^50^ were excluded from the analysis. The remaining reads were aligned to the human genome hg38 and assigned to the GENCODE Human release 32 reference using STAR. UMI suppression was performed using UMItools v1.1.2^51^. Downsampling of datasets was performed using seqtk (https://github.com/lh3/seqtk). The definition of representative genes was based on MANE select v.1.0^52^.

For the standard Ribo-Seq, the A-site offsets were determined to be 15 for footprints of 20-22 and 24-31 nt and 16 for footprints of 23 and 32 nt, based on the metagene analysis of the 5’-end of footprints. For Thor-Ribo-Seq, the A-site offsets were defined as 15 for footprints of 21-22 and 27-31 nt and 16 for footprints of 23 and 32-33 nt. Ribosome occupancies were defined as the number of reads at given codons normalized by the average reads per codon on each CDS. Transcripts with 1 or more reads per codon on average in all samples were considered.

### Resource availability

Standard Ribo-Seq and Thor-Ribo-Seq data (GEO: GSE222195) were deposited in the National Center for Biotechnology Information (NCBI) database. Further information and requests for resources, reagents, and scripts should be directed to and will be fulfilled by the Lead Contact, Shintaro Iwasaki.

---

**Figure S1.**
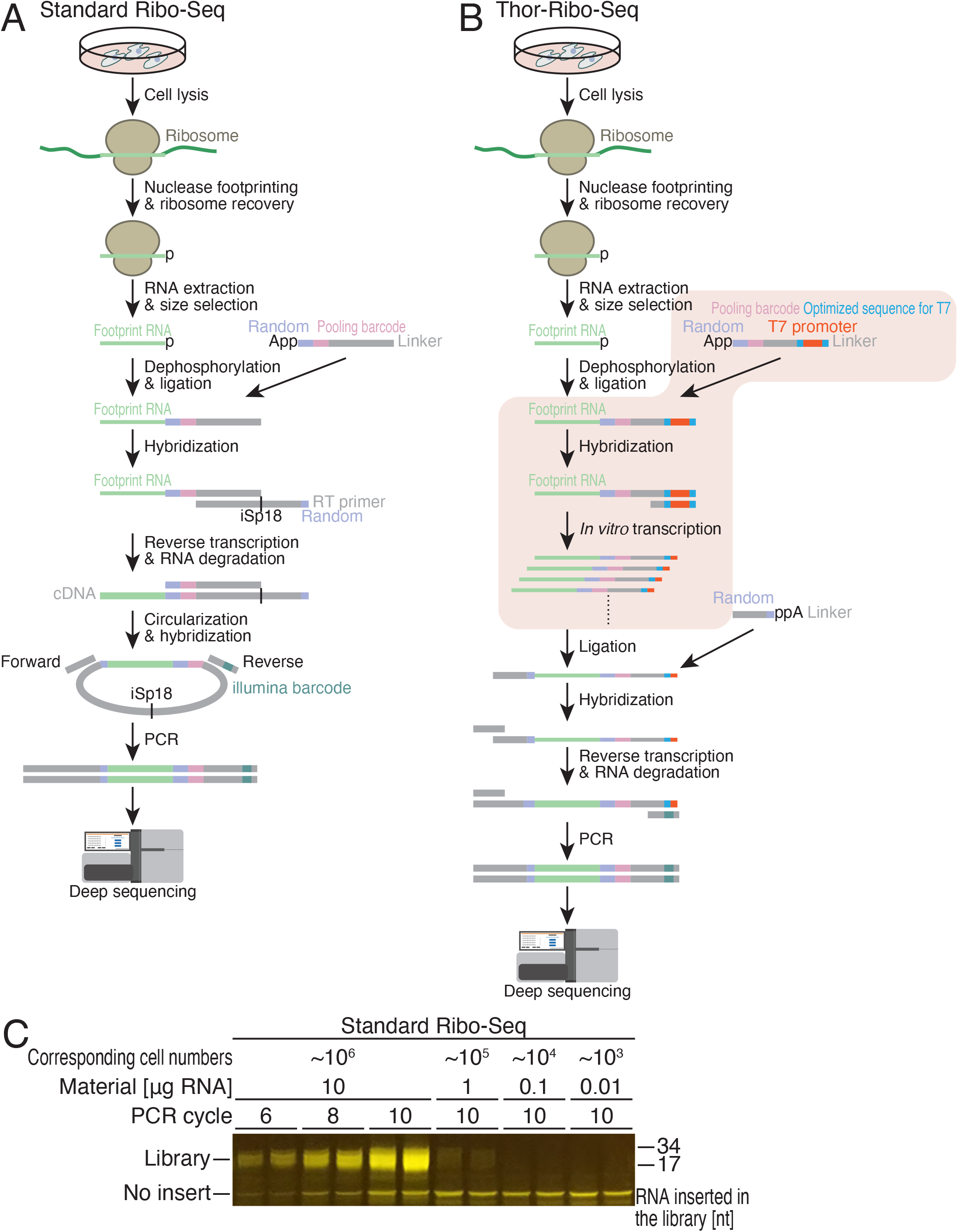
The limitation of the standard Ribo-Seq strategy for low inputs. Related to Figure 1. (A and B) Step-by-step overviews of library preparation schemes for standard Ribo-Seq (A) and Thor-Ribo-Seq (B). Random sequences were used as UMI. (C) A gel image of the PCR-amplified DNA library prepared by the standard Ribo-Seq method with titrated amounts of initial materials.

**Figure S2.**
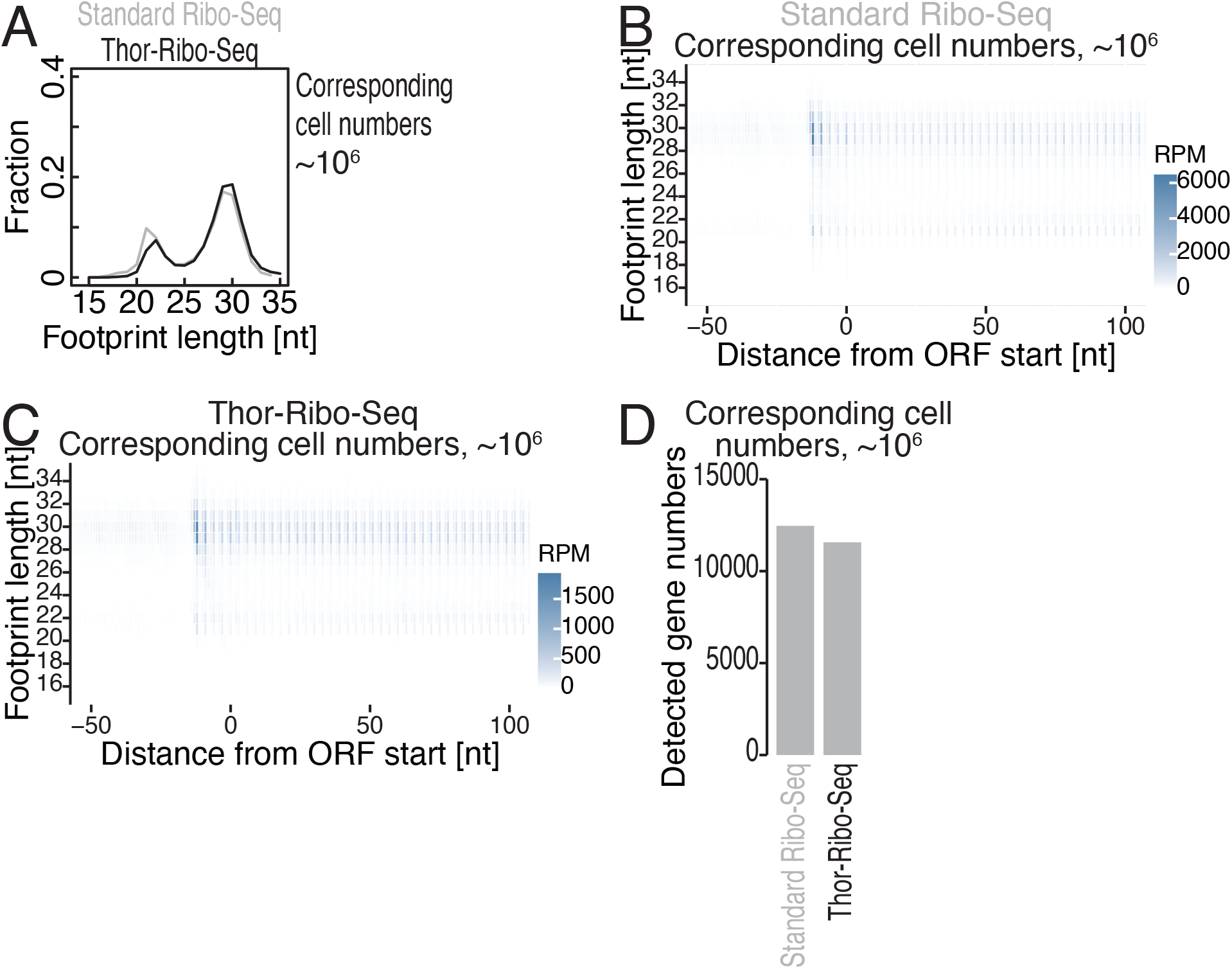
Hallmarks of ribosome footprints obtained by Thor-Ribo-Seq with a normal input. Related to Figure 1. (A) The distribution of footprint length for standard Ribo-Seq and Thor-Ribo-Seq with normal inputs. (B and C) Metagene plots of the 5’ end position of ribosome footprints around start codons in the indicated experiments. X axis, the position relative to the start codon (0 as the first nucleotide of the start codon); Y axis, footprint length; color scale, read abundance. RPM, reads per million reads. (D) Detected gene numbers (3 reads or more) in 4 million footprint reads in the indicated experiments. Representative transcripts in each gene defined in the MANE Select (Ensembl) were used.

**Figure S3.**
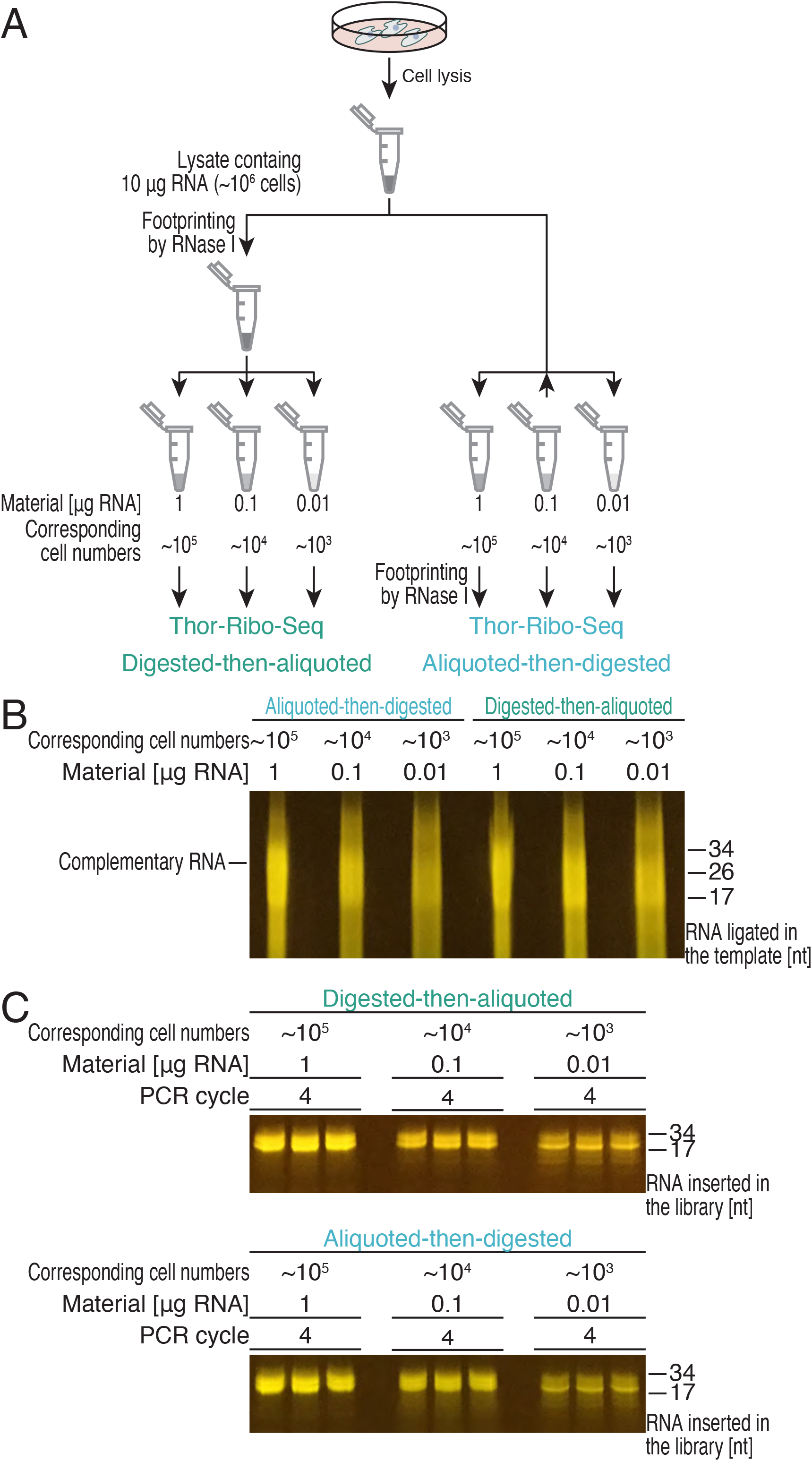
Library preparation with Thor-Ribo-Seq method from low inputs. Related to Figure 1. (A) Schematic representation of experimental designs to test the potential of Thor-Ribo-Seq. In the “digested-then-aliquoted” experiments, lysates were treated with RNase I and then divided into different amounts. In the “aliquoted-then-digested” experiments, lysates were first titrated and then digested with RNase I. (B) A gel image of *in vitro* transcription-amplified complementary RNA to the ribosome footprints. (C) A gel image of the PCR-amplified DNA library prepared by the Thor-Ribo-Seq method with titrated amounts of initial materials.

**Figure S4.**
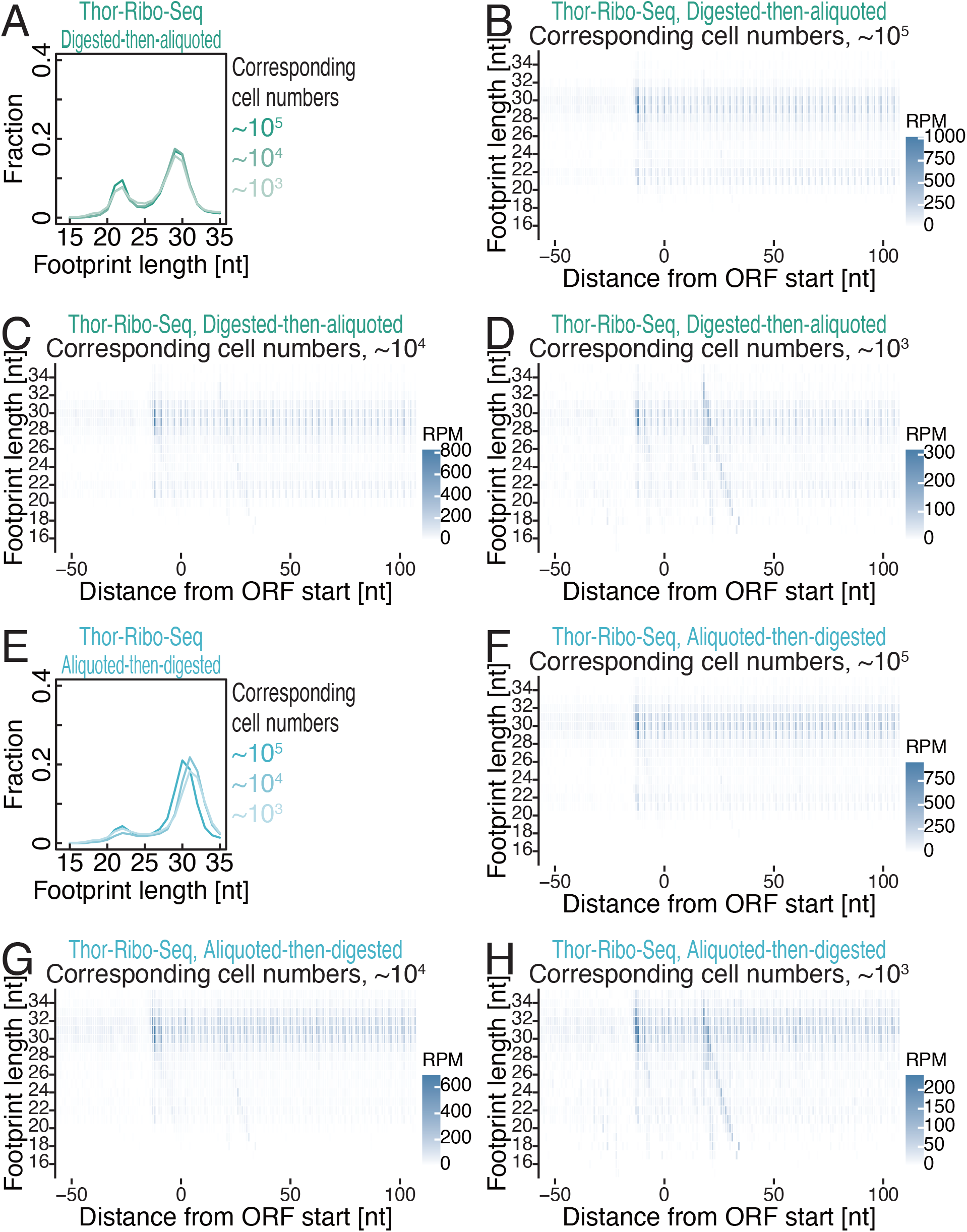
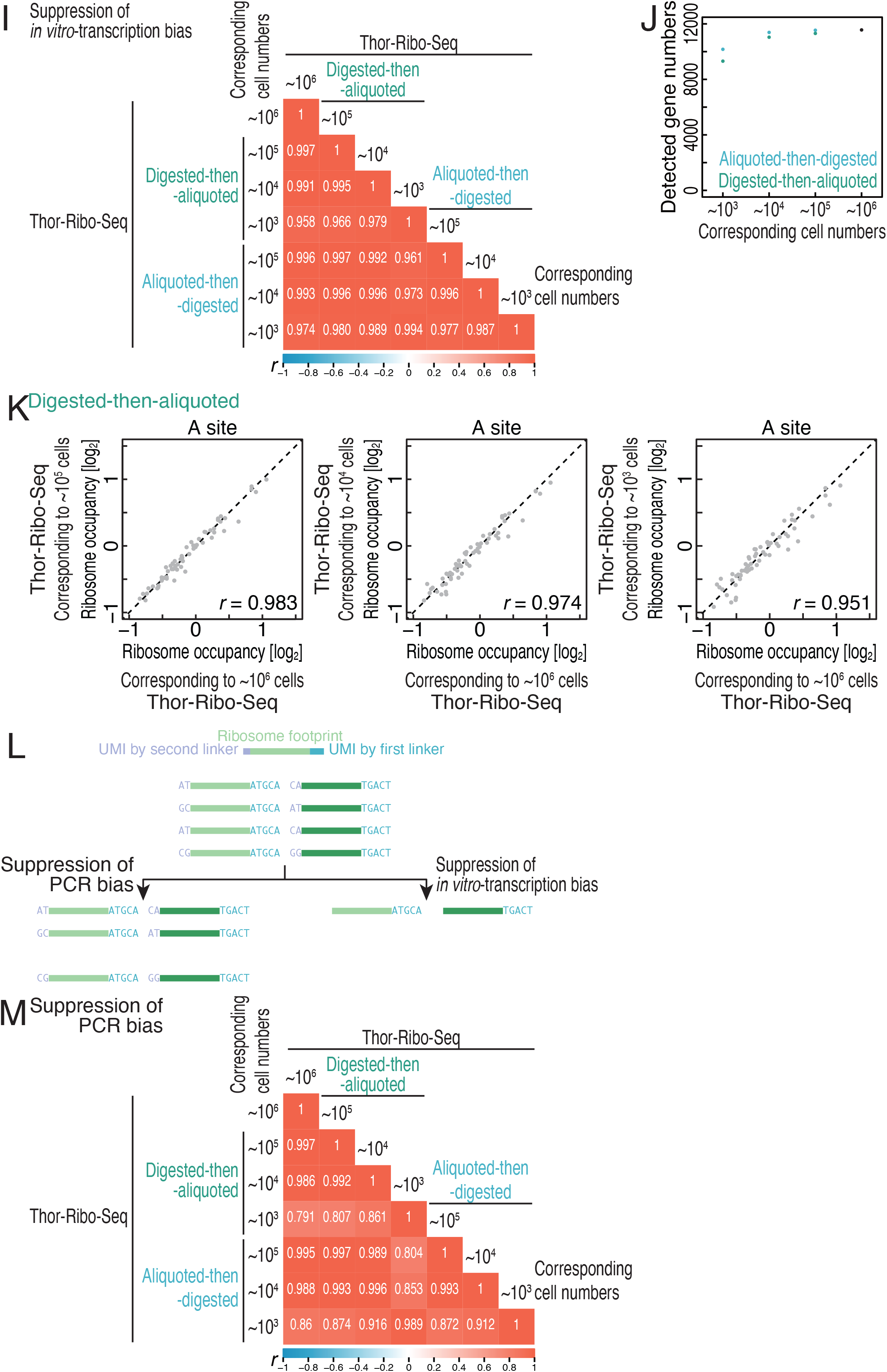
Investigation for biases in Thor-Ribo-Seq originated from low inputs. Related to Figure 1. (A and E) The distribution of footprint length for Thor-Ribo-Seq with “digested-then-aliquoted” conditions (A) and that with “aliquoted-then-digested” conditions (E). (B-D and F-H) Metagene plots of the 5’ end position of ribosome footprints around start codons in the indicated experiments. X axis, the position relative to the start codon (0 as the first nucleotide of the start codon); Y axis, footprint length; color scale, read abundance. RPM, reads per million reads. (I) Pearson’s correlation coefficient (*r*) among the indicated experiments. Read duplications generated by *in vitro* transcription were suppressed by the UMI in the first linker. The *r* value scales are shown at the color bars. (J) Detected gene numbers (3 reads or more) in 4 million footprint reads in the indicated experiments. Representative transcripts in each gene defined in the MANE Select (Ensembl) were used. (K) Correspondence of averaged ribosome occupancy on A-site codon sequences across Thor-Ribo-Seq (digested-then-aliquoted) with low material inputs. *r*, Pearson’s correlation coefficient. (l) Schematic representation of UMI-mediated duplicated read suppression. (M) Pearson’s correlation coefficient (*r*) among the indicated experiments. Read duplications generated by PCR were suppressed by the UMI in the first linker and the second linker. The *r* value scales are shown at the color bars.

